# Mapping Allosteric Rewiring in Viral RNA: Sequence-Encoded Control of Protein Binding Mechanisms

**DOI:** 10.1101/2025.11.01.685830

**Authors:** D Samant, A Chakraborty, A Sinha, R Sarkar, S Roy

**Affiliations:** Department of Chemical Sciences, Indian Institute of Science Education and Research Kolkata, West Bengal 741246

**Keywords:** HIV-1 TAR-Tat, HIV-2 TAR-Tat, Single Point Mutation, RNA Allostery

## Abstract

RNA recognition by proteins is governed not only by static structure but also by allostery encoded within non-local dynamic motifs. In this study, we systematically identify allosteric communication hubs in RNA and map multiple residue-connected pathways, revealing how these networks are rewired upon mutation and protein binding. To capture these effects under physiological salt conditions, we performed tens of microseconds of atomistic and steered molecular dynamics simulations and computed binding free energies for Tat–TAR complexes across three immunodeficiency virus variants, BIV, HIV-1, and HIV-2. Allosterically coupled sites were identified using contact-based principal component analysis, and communication pathways were traced through an extended graph-network algorithm—the first such application to RNA systems. Two distant motifs—the bulge and the apical loop—emerge as allosteric switches and information hubs: the bulge engages Tat, while the loop interacts with another protein partner, CycT1, both essential for transcriptional activation and antiviral targeting. We find that HIV-2 TAR, with strong loop–bulge coupling and high self-integrity, favour conformational selection and exhibits lower Tat-binding affinity. In contrast, a single C24 insertion in HIV-1 TAR reconfigures communication pathways, enabling an induced-fit mechanism with enhanced affinity. The study not only elucidates an allosteric rewiring between the loop and bulge but also highlights how this communication is dynamically reconfigured upon protein binding. Tat association at the bulge reorganizes and reorients loop residues, thereby promoting the subsequent recruitment of CycT1. This work overall underscores how sequence (even a single mutation) encoded RNA allostery can modulate not only a protein’s binding mechanism and affinity but also influence downstream molecular events within transcriptional signalling cascades.

**STATEMENT OF SIGNIFICANCE:** In vitro selection strategies such as SELEX have long underscored the therapeutic relevance of the TAR RNA–Tat protein interface, yet they remain experimentally demanding and often overlook how subtle sequence variations reshape molecular recognition and RNA-protein binding affinity. By integrating atomistic and free-energy simulations with a graph-network-based framework, the study reveals that even a single-nucleotide change can rewire RNA’s allosteric communication pathways, switching between conformational selection and induced-fit mechanisms and alter binding affinity. These findings bridge molecular dynamics with experimental selection, offering a predictive, thermodynamics-based framework to rationalize sequence-structure-function relationships and guide the rational design of high-affinity RNA-protein complexes and next-generation RNA therapeutics.

## INTRODUCTION

RNA-protein complexes are fundamental molecular machines that orchestrate diverse cellular processes, from transcription and translation to various intermediate stages of gene regulation across all domains of life(1–4) . Their versatility arises from the synergistic interplay between RNA and protein, whose intrinsic dynamism enables a multitude of interfacial interactions and diverse binding modes.

A prototypical example of this interface complexity is the TAR-Tat interaction in lentiviruses such as HIV-1, HIV-2, and BIV (Human Immunodeficiency Virus: pathogenic; Bovine Immunodeficiency Virus: non-pathogenic(5)). The structured RNA element TAR, located in the 5′ untranslated region of nascent viral transcripts, recruits the viral Tat protein to initiate transcriptional elongation(6–8) by assembling host cofactors such as P-TEFb (Cyclin-T1 and CDK9)(9, 10) . Given its central role in hijacking host transcription machinery, the TAR-Tat complex has long been considered a promising therapeutic target.

Within this paradigm, in vitro selection strategies such as SELEX (Systematic Evolution of Ligands by Exponential Enrichment) have been extensively used to engineer aptamers that mimic TAR and disrupt its high-affinity binding to Tat(11, 12). Large combinatorial libraries of randomized sequences are typically screened to identify aptamers with binding specificity. In 1998, SELEX applied to ∼10^13^ RNA variants yielded TAR aptamers that retained the Tat-binding domain while maintaining specificity comparable to wild-type TAR(13). Later, Yamamoto et al. reported a 37-nt hairpin RNA aptamer (RNA^Tat), derived from ∼10^14^ randomized 120-nt sequences, which displayed two-fold higher Tat affinity than wild-type TAR and functioned as an effective molecular recognition element even in the presence of excess of wild type-TAR(14). While the above experiments underscore the therapeutic potential of targeting TAR-Tat, SELEX approaches remain experimentally demanding, relying on vast combinatorial libraries and iterative enrichment cycles to identify high-affinity binders. Moreover, subtle natural variations among viral TAR RNAs can fundamentally alter recognition pathways, raising questions about the generalizability of such aptamers. Here, complementary computational strategies can add value by mapping competent RNA motifs, they can uncover mechanistic determinants of affinity and highlight sequence regions most likely to function as natural “switching knobs.” Such insight can help reduce the experimental burden by narrowing sequence space prior to SELEX, thereby guiding more targeted experimental designs.

Structurally, TAR RNA comprises two helical stems connected by an asymmetric U-rich bulge and capped by an apical loop(15–18) . Tat recognition is primarily mediated through its arginine-rich motif (ARM) binding into the bulge pocket(15, 19, 20) . NMR studies revealed that U23–A27:U38 base triple formation generates a unique recognition pocket near the bulge(21, 22). Follow up NMR studies showed significant apo-state rearrangements upon Tat binding, especially reflected in the bulge(23). Mutational studies further demonstrated that bulge size and geometry strongly modulate binding affinity(20, 24) . Interestingly, the natural TAR systems are also observed to tune bulge feature. HIV-2 TAR notably lacks the C24 nucleotide present in HIV-1 TAR(25, 26). Although Weeks and Crothers showed that even a two-base bulge suffices for Tat binding, underscoring the importance of geometry over absolute bulge size(24) , experimental Scatchard analyses by Karn et al. confirmed that bulge flexibility enhances Tat affinity(27). Based on these experiments, Jerningen and coworkers employed a lattice model to show how larger, more flexible bulges may increase conformational adaptability facilitating protein accommodation(28). While these findings point to the correlation between large bulge entropy of the apo-RNA with RNA-protein binding affinity, but affinity cannot be fully explained by entropy alone without detailed free-energetic characterization.

Beyond the bulge, the TAR loop also plays an essential role in transactivation(29, 30). Dingwall et al.(15, 31) proposed the loop as a potential cofactor recruitment site(30). Neenhold et al.(32) observed that Tat binding at the bulge perturbs loop dynamics, priming it for additional host factor engagement. Mutational studies by Berkhout et al. demonstrated that instead of individual the spatial relationships between loop and bulge modulate transcriptional efficiency(33). Dethoff et al. revealed long-range bulge-loop coupling in apo TAR RNA, suggesting cooperative conformational signaling(34). In fact, in one of our prior studies on BIV TAR RNA, we showed that anti-correlated loop-bulge fluctuations establish a favorable recognition landscape that promotes high-affinity Tat binding(35). In contrast, HIV-2 TAR requires Tat-induced loop activation and additional cofactors (Cyclin-T1) to stabilize its RNA structure(36), pointing to distal allosteric cross-talk and a more complex multicomponent assembly mechanism. Nonetheless, the microscopic mechanism of this inter-motif crosstalk and its effects on binding affinity remains unclear.

These comparative insights underscore the functional divergence among different TAR RNA sequences despite structural similarity. Subtle sequence variations can remodel RNA-protein recognition strategies, altering both binding landscapes and functional outcomes. A particularly intriguing case is HIV-1 TAR’s additional cytosine (C24) in its bulge, absent in HIV-2. This single-nucleotide difference is linked to stronger Tat binding affinity (HIV-1: K_d_ = 12 nM; HIV-2: K_d_ = 30 nM)(27). This raises fundamental questions: can the additional C24 in HIV-1 TAR act as a molecular switch that modulates bulge plasticity and rewires bulge–loop allostery to enhance Tat affinity? If so, what microscopic mechanisms allow a single nucleotide substitution to reshape allosteric networks and fine-tune the thermodynamic and structural landscape of RNA-protein recognition? These issues remain unresolved. Addressing them is not only central to understanding the evolutionary strategies of viral regulatory RNAs but also to uncovering generalizable design principles of RNA allostery. In particular, identifying how local sequence edits function as intrinsic ‘control knobs’ of affinity and binding mode could establish a rational framework for RNA engineering-one that complements experimental approaches and informs the design of therapeutics targeting TAR-Tat and other RNA-protein complexes.

Continuing from our early study(36) to elucidate how two distal active sites, the loop and bulge, in TAR RNAs from immunodeficiency viruses govern intrinsic structural properties, regulate overall integrity, and thereby control modes of protein accommodation, at the beginning, we compared the structural and dynamical behavior of two pathogenically distinct(5) but structurally analogous TAR RNAs. Both share a stem-loop-bulge architecture: one from Bovine Immunodeficiency Virus (BIV PDB id:1BIV (37)) and the other from Human Immunodeficiency Virus-2 (HIV-2(38), PDB id: 6MCE (38)). For this comparison, we performed all-atom molecular dynamics equilibrium simulations in explicit solvent environment, maintaining physiological ion concentration, as detailed in the method section.

## METHODS

### System Setup

The initial atomic coordinates of the TAR-Tat complex corresponding to HIV-2 were directly obtained from the Protein Data Bank (PDB ID: 6MCE (38)), representing the high-resolution solution NMR structures reported by Pharm et al. Since the full-length TAR-Tat complex structure for HIV-1 has not yet been experimentally resolved, it was merely constructed by manually inserting a C24 nucleotide at the bulge region and removing the terminal G:C base pair from the HIV-2 framework using UCSF Chimera software. The resulting TAR-Tat complexes are illustrated in the **Figure S1**. To examine the apo RNA conformational dynamics, the HIV-2 TAR structure was derived by eliminating Tat peptide coordinates from the initial TAR-Tat complex, as no free HIV-2 TAR RNA is currently available. In contrast, HIV-1 apo TAR solution NMR structure was sourced from Protein Data Bank (PDB ID: 1ANR (39)), as reported by Aboul-ela et al..

### Simulation protocols

All the atomistic simulations were performed using GROMACS 2018.3 package (40). System-specific topologies were generated employing the Amber 99 force field(41) with parmbsc0(42) and chiOL3(43) refinements. Each system was centered in a cubic box with dimensions of 12 nm per side and solvated using the TIP3P water model. To ensure charge neutrality and maintain a physiological ionic strength, sodium and chloride ions were added to achieve desired salt concentration. The ion and solvent counts have been detailed in **Table S1** of the Supporting Information.

To eliminate steric clashes and optimize geometry, following system preparation, energy minimization was performed using steepest descent algorithm. During the initial equilibration phase, positional restraints with a force constant of 1000 kJ/mol.nm^2^ were imposed on the systems, while the ions were kept frozen under NVT conditions for 3 ns to ensure proper solvation. Subsequently, the ionic restraints were released and the systems were further equilibrated for an additional 3 ns. To gradually relieve positional constraints, the restraint force constants were systematically reduced from 1000 to 10 kJ/mol.nm^2^ in intervals of 3 ns. This was followed by complete removal of all restraints during 1 ns equilibration under the NPT ensemble. Afterward, unrestrained equilibrium production runs were carried out under NVT ensemble.

All the simulations employed leapfrog integrator with a 2 fs time step. Temperature was maintained at 300 K using the Nosé-Hoover thermostat(44, 45) with a coupling time constant of 0.5 ps. For all pressure-controlled (NPT) equilibration steps, pressure was maintained at 1 bar using Parrinello-Rahman barostat(46) with a time constant of 0.5 ps. Periodic boundary conditions were applied in all directions. The neighbor list was updated every 10 steps using a grid-based algorithm with a short-range cutoff of 1.0 nm. Long-range electrostatic interactions were computed using the Particle Mesh Ewald (PME)(47) method with a Fourier grid spacing of 0.12 nm and fourth-order interpolation. All bond constraints were enforced using LINCS algorithm(48).

### Bulk Ionic Strength Estimation

Time-averaged bulk salt concentrations were computed by evaluating the time-averaged ratio of the number of bulk ions to the number of bulk water molecules, scaled by the molarity of pure water. Based on the radial distribution function shown in **Figure S2**, all species located beyond 2 nm from the RNA were defined as bulk and only those were considered in the calculation (35).

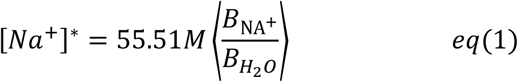

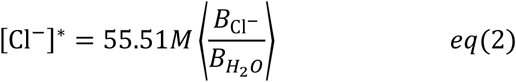

To account for potential electrostatic imbalances inherent to molecular dynamics simulations, these raw concentrations were corrected using a charge-neutrality correction relation as:

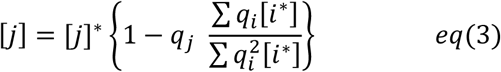

Specifically, the corrected concentration [j] of an ion obtained from its raw concentration [j]* using the relation adjusts for the net charge contributions of all ionic species present in the system. This ensures that the final concentrations reflect a physically meaningful, charge balanced ionic environment. Following this correction, the bulk concentration of NaCl in both systems was found to be approximately 100±10mM, consistent with physiological ionic strength. Detailed values of the ion concentration are provided in **Tables S2-S3**.

### Constant velocity steered MD

To investigate the pathway of dissociation of protein from the RNA groove, we employed constant velocity steered molecular dynamics simulation (cv-SMD (49)). This technique introduces a time-dependent external force along a predefined reaction coordinate, enabling the system to surmount the energy barrier involved in the process of dissociation. By applying a harmonic bias potential, the protein was gradually pulled away from the RNA at a uniform velocity, effectively facilitating the dissociation event. Unlike conventional molecular dynamics, which may be limited by the thermal energy available at room temperature, cv-SMD enables access to otherwise inaccessible unbinding intermediates. The simulations were conducted by defining centre-of-mass distance between the TAR RNA and the Tat peptide as the reaction coordinate using GROMACS 2018.3 package (40). A force constant of 500 kJ/mol.nm^2^ and a pulling velocity of 0.0001 nm/ps were employed to ensure controlled and progressive unbinding.

### Free energy calculation method

To compute the free energy profile, umbrella sampling(50) method was performed using the centre-of-mass distance between the RNA and the protein as reaction coordinate. Initial configurations for each window were extracted from the cv-SMD trajectories. These configurations were selected by ensuring an adequate overlap between adjacent windows, thereby enhancing phase space coverage. Each window was simulated for 10ns under NVT ensemble at 300K, using a harmonic force constant of 500 kJ/mol.nm^2^ and a pulling velocity of 0 nm/ps.

Temperature regulation was maintained via the Nosé-Hoover thermostat (51, 52). The overlapping of windows ensured continuous sampling along the reaction coordinate (42 windows for HIV-2 and HIV-1, the windows overlapping has been shown in **Figure S3**). The resulting biased simulations were then analyzed using Weighted Histogram Analysis Method (WHAM(53)), which combines the individual potential of mean force (PMF) profiles to generate an unbiased PMF profile, upon which an entropy correction resulted in free energy landscape. The statistical error was calculated from the Bayesian Bootstrapping(54).

The free energy profiles obtained here represent only the barrier associated with dissociation of RNA and Protein, as characterized by their centre-of-mass (COM) distance, chosen as an 1D order parameter. While COM distance captures the global separation appropriately, alone it may lack capturing the right configurational states along the pathway. The resulting free energy analyses compare a loop-guided recognition pathway in different variants rather than an absolute description of the whole dissociation mechanism, which might be governed by any/ combination of alternative suitable order parameter/s. Besides, the conformational transitions along the pathway inherently depends on the explicit solvent conditions of simulations.

The detailed durations of both the biased and unbiased simulations are presented in **Table S4-S6**.

### Contact-Based Principal Component Analysis

To identify long-range dynamic coupling between the bulge and loop regions in the apo states of the RNAs, we performed contact-based principal component analysis (contact-PCA) using equilibrium simulation trajectories. This approach captures the principal modes of contact fluctuations, thereby revealing domain-level coupling within the RNA structure. Minimum distances between all heavy atoms were computed for every unique pair of residues. For HIV-2 apo RNA with 30 residues, having 435 unique pairs; for the HIV-1 apo RNA, which consists of 29 residues, 406 unique residue pairs were obtained. A binary contact map was generated for each frame of the trajectory using a distance cutoff of 0.5 nm (i.e., a contact was assigned a value of 1 if the minimum distance ≤ 0.5 nm, and 0 otherwise). Next, a covariance matrix was constructed from the time series of binary contact data. This matrix was diagonalized, and principal component analysis was performed via eigenvalue decomposition. The first principal component (PC1), corresponding to the largest eigenvalue, was considered for the analysis. The absolute values of the feature vector components (i.e., the unique residue–residue contact pairs) of PC1 were calculated, representing the contribution (or weight) of each residue pair to the dominant contact fluctuation mode. The top five residue pairs with the highest absolute weights were extracted and mapped onto the RNA structure to interpret the key contacts mediating dynamic coupling between the bulge and loop domains.

### Contact Probability Map

The RNA-protein binding interface was further examined by the evaluating the contact probability maps. To construct the same for RNA-protein interactions at the residue level resolution, first of all an all-atom contact map by considering only the heavy atoms of both the TAR RNA and Tat peptides was generated. This atomistic contact map was subsequently converted to residue-level resolution using the following algorithm.

The contact cut-off distance was decided based on the radial distribution function of all heavy atoms of RNA surrounding the heavy atoms of protein (**Figure S4**). A cut-off of 4.3 A° was identified as the threshold for defining atom pairs in contact. At each simulation frame, we constructed a binary contact matrix of dimensions [RNA atom number × Protein atom number], where each matrix element was assigned a value of 1 if the corresponding atom pair was within the cutoff distance and 0 otherwise.

Summing these matrices over all frames yielded the total number of contact occurrences for each atom pair across the trajectory. We identified the atom pair with the highest frequency and normalized all contact frequencies by this maximum value or total number of equilibrium frames, resulting in a normalized all-atom contact map with values ranging from 0 to 1.

To convert this data to residue-level resolution, each atom was mapped to its corresponding residue index. For any given RNA-protein residue pair, multiple atom-atom interactions may exist. Among these, the atom pair with the highest normalized frequency was selected to represent the contact strength for that particular residue-residue interaction. While this residue level representation abstracts away some of the fine-grained details captured at the atomistic level, it provides a clear and interpretable overview of the contact sensitivity between specific RNA nucleotides and protein residues.

### Graph-Network Construction to Map Allosteric Path

Furthermore, to delineate the plausible pathway of allosteric channel within TAR RNA, we have constructed graph-based networks, first-time for 3D RNA structures. Dynamical network analysis is a correlation-based methodology that leverages molecular dynamics (MD) simulations to elucidate pathways of allosteric communication within biomolecules(55, 56). In this framework, the system is represented as a graph, where nodes correspond to structural elements such as amino acid residues and nucleotides, and edges represent their interactions.

At the initial stage, the network is constructed at the heavy-atom level for RNA in HIV-2 and HIV-1 apo, and for both protein and RNA in the HIV-2 and HIV-1 complex. An edge is defined between two nodes if the corresponding heavy atoms (non-hydrogen) are in contact, within 5 Å for a minimum of 45% of equilibrium frames of trajectory. These parameters are selected to capture pairs that display significant correlations, even if their overall contact frequency across the trajectory is relatively modest. Unlike prior approaches(57) that exclude sequential neighbors to avoid trivial connections, this study retains adjacent residues to allow exploration of the full spectrum of potential pathways, including suboptimal routes, within the weighted network.

The dynamical networks were constructed from the ∼1µs simulation data, with trajectory frames sampled every 100ps time step. The resulting graphs for RNA in HIV-2 and HIV-1 apo, and for both protein and RNA in the HIV-2 and HIV-1 complex, are weighted, with the edge weight *w_ij_* between nodes i and j defined as:

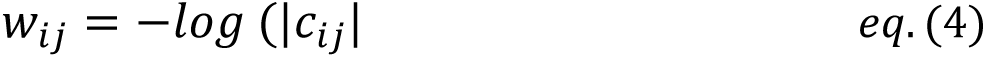

where *c_ij_* is the cross-correlation value between node i and j. To focus on the most significant contributors to flow the information from the source (bulge) to the sink (loop) in HIV-2 (apo and complex) and HIV-1 (apo and complex), the top 30 heavy atoms exhibiting the strongest correlations with their network connections were extracted and the shortest signalling pathways were identified using Dijkstra’s algorithm(58), as implemented in the NetworkX Python library(59). This framework naturally interprets weights in terms of information transfer probabilities, enabling a probabilistic view of path lengths. Both positive and negative correlations are incorporated symmetrically. For HIV-2 (apo and complex) and HIV-1 (apo and complex), residue- and nucleotide-level representations were then obtained by projecting atom-level correlations onto their corresponding residues or nucleotides, with each treated as a single node.

Visualization and analysis of communication pathways were carried out by mapping network properties to graphical attributes: node size and color represent the recurrence frequency of residues/nucleotides across computed pathways, where highly recurrent residues/nucleotides appear as larger, red nodes, and less frequent contributors appear smaller and paler but may still play nontrivial roles. Edge thickness is proportional to the number of highly correlated atoms interacting between two residues/nucleotides, such that frequently interacting correlated residue/nucleotide pairs (lower weights) are shown as thicker edges. This residue-level network representation provides a systematic means to evaluate the contributions of individual nucleotides to allosteric signalling and offers a physically interpretable model for how signals propagate in HIV-2 (apo and complex) and HIV-1 (apo and complex) systems.

The detailed overviews of the rest of the trajectory-based analyses including calculations of RMSD, RMSF, intra- and inter-chain interaction energies and covariance has been illustrated in Supporting Information.

## RESULTS AND DISCUSSION

### A Single Point Mutation in Active Site Bulge of Apo HIV TAR RNA Alters Distant Loop Dynamics: Allosteric Onset

Apo TAR RNAs, being intrinsically flexible, often exhibit correlated fluctuations across spatially separated domains. Early experimental studies and our early works on ***BIV TAR RNA **(PDB id:1BIV)** (37) have indicated that the coupling between two such distant domains-loop and bulge **(Figure S5**) in TAR RNA might be relevant to Tat binding (35). However, how their coupling may interfere the binding and what are the specific features of loop and bulge are determinant factors-these critical questions we ask in this study. At this point, from early literatures, we surmise several critical loop-bulge features of apo-RNA that likely govern its protein affinity(36): (i) bulge length, (ii) long-range loop–bulge proximity, (iii) dynamic correlation between loop and bulge fluctuations, and (iv) RNA’s overall self-integrity. These features, however, can be interdependent or independent. For example, increasing bulge length may enhance RNA flexibility but does not necessarily ensure loop-bulge proximity or correlated fluctuations. Similarly, loop-bulge proximity does not inherently imply dynamic coupling. To rigorously test the role of bulge length and associated features, instead of comparing more distantly related TAR RNAs, we turned to HIV-1 TAR, which differs from HIV-2 by only a single nucleotide insertion: the cytosine at position 24 (C24) (**Figure 1A-B**). This subtle variation allowed us to investigate both apo and complex states while keeping the partner protein (Tat-1) identical. Using the solution NMR structure of HIV-1 TAR (PDB id: 1ANR(39)), we simulated its apo dynamics and compared them with HIV-2. RMSF analysis (**Figure 1C**) revealed that although the nucleotide addition occurs at the bulge, it induces enhanced fluctuations at distal loop residues such as G32 in HIV-1, a feature absents in HIV-2. To probe this mutation-induced redistribution of dynamics, we calculated fluctuation-based covariance matrices **(Figure S5**), which capture correlated and anti-correlated positional variations across residues. HIV-2 apo TAR displayed strong loop-bulge anti-correlation, consistent with tightly integrated domain dynamics. In contrast, HIV-1 apo TAR exhibited weaker and more delocalized correlations, suggesting less integrated long-range dynamics-consistent with differences observed in interaction energy (**Figure S1**).

**Figure 1:**
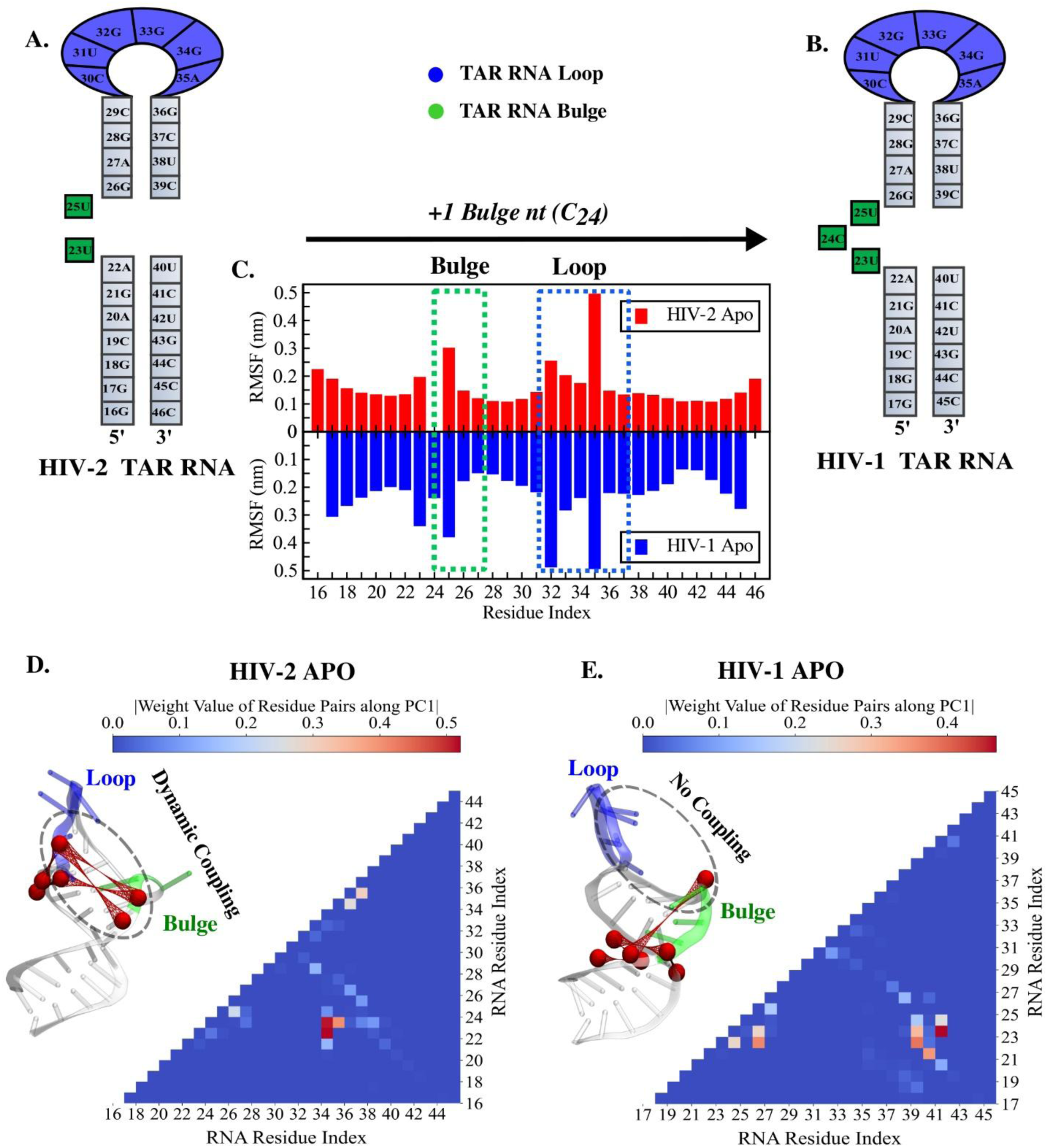
Effect of a single point mutation on the apo state loop-bulge dynamics in HIV-TAR RNAs. **(A) and (B)** represent the secondary structures of HIV-2 and HIV-1 TAR RNAs respectively, connected via a single point mutation (C24) at bulge. **(C)** compares residue wise time-averaged root mean square fluctuation (RMSF) between HIV-2 and HIV-1 apo TAR. **(D) and (E)** illustrate the dynamic coupling between the bulge and loop regions in the HIV-2 and HIV-1 apo RNA structures respectively, as captured by contact-PCA analysis. The top five residue–residue contact pairs (ranked by their weight values) are projected onto the RNA structures. Red van der Waals spheres represent the phosphate beads of the RNA, and the connections between residues are shown using red wireframes. The structures were visualized using VMD.

To further resolve the specific contacts underlying these inter-domain couplings, we applied contact-based principal component analysis (contact-PCA) at residue-level resolution. Mapping the top five residue pairs with the largest weights onto RNA structures revealed the key long-range contacts that mediate loop-bulge cross-talk in HIV-2 apo TAR (**Figure 1D**). Strikingly, HIV-1 TAR lacked such robust long-range coupling (**Figure 1E**), underscoring the disruptive impact of the single-nucleotide insertion on allosteric connectivity.

### Binding Free Energy Uncovers Different Modes of Protein Accommodation in RNA Groove

A single-point mutation, namely the addition of C24 in HIV-1 TAR, indeed markedly alters RNA dynamics and rewires loop-bulge coupling (**Figure 1**). Yet, how this dynamic rewiring in HIV-1 TAR (3-nt bulge) influences the Tat encounter process and binding thermodynamics compared to HIV-2 TAR (2-nt bulge) required quantification. To address this, we first quantified intra-chain interaction energies in apo TAR RNAs and then compared inter-chain interactions in complexes **(Figure S1)**. In the apo state, BIV TAR shows lower self-integrity than HIV-2, enabling stronger protein mixing and yielding a more stable RNA-protein complex. Conversely, HIV-2 TAR maintains higher apo-state self-integrity, correlating with weaker protein interactions and comparatively less stable complexes. HIV-1 TAR, in contrast, exhibits an intermediate balance of flexibility (**Figure S5-S6**) **and affinity, in comparison to BIV and HIV-2 cases (Figure S1, Figure S3)**.

Now to evaluate and compare the thermodynamics, we employed umbrella sampling to compute Free Energy profiles for TAR–Tat dissociation. These profiles, averaged over multiple pulling pathways derived from steered molecular dynamics (SMD) simulations along the RNA-protein centre-of-mass (COM) distance reveal a unique trend: HIV-2 exhibits a markedly lower dissociation barrier (**Figure 2A**) than HIV-1 (**Figure 2B**). These results mirror interaction energy patterns **(Figure S1**) and align with earlier experimental measurements of dissociation constants (K_d_)(27), reinforcing the principle that less self-integrated TAR RNAs form stronger adducts with Tat.

**Figure 2:**
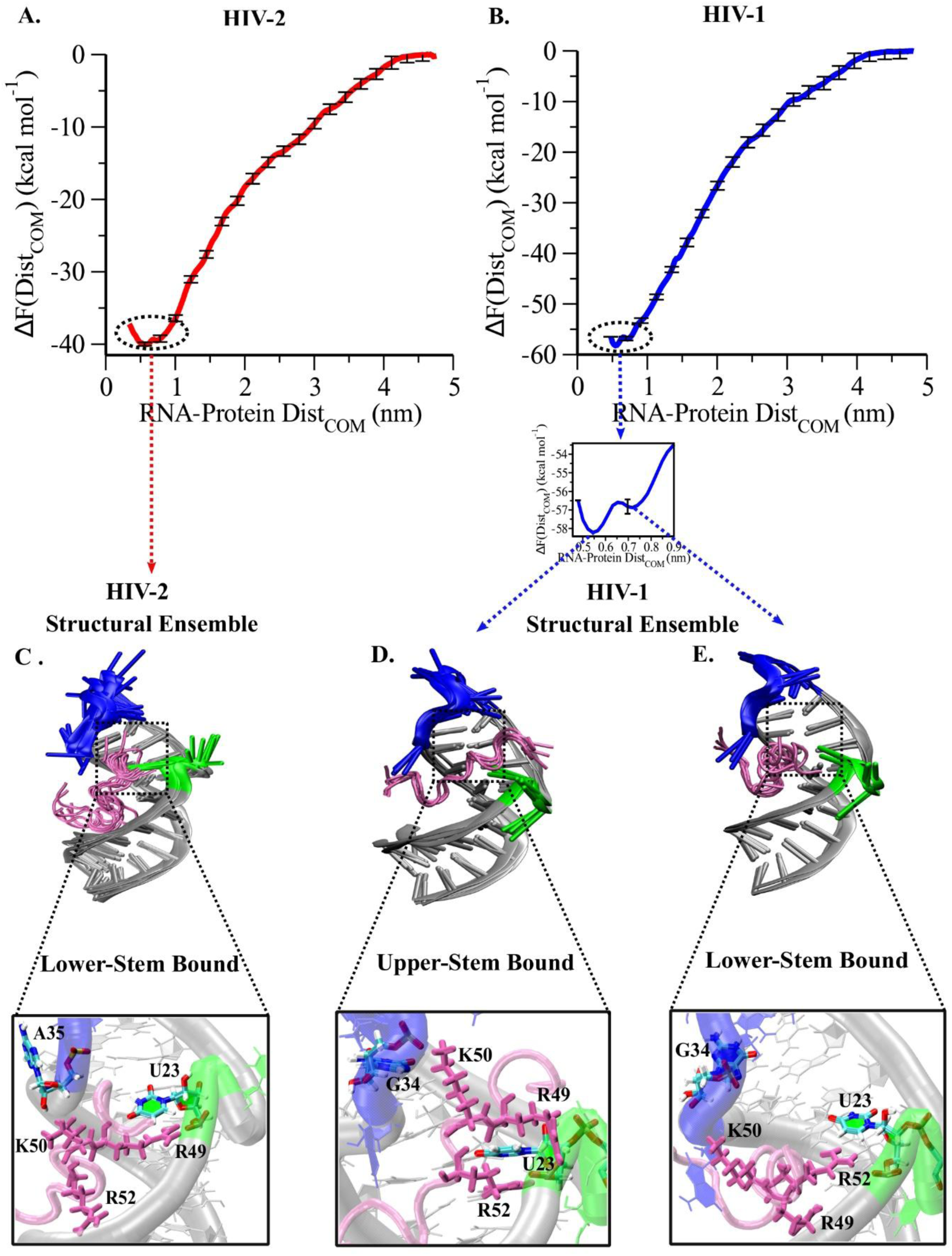
Effect of a point mutation in TAR affecting modes of Tat accommodation and affinity. **(A)** and **(B)** depict the free energy profiles obtained upon dissociating the Tat protein from TAR RNA’s major groove along their centre of mass distances for HIV-2 (red) and HIV-1 (blue) respectively. The profiles have been scaled referring the bulk at zero. **(C)** represents seven distinct snapshots made out of equilibrium trajectories of HIV-2 TAR-Tat complex to show the heterogeneous interactions phase space along with their microscopic interactions, **(D)** and **(E)** represent structural ensemble for HIV-1 TAR-Tat complexes and their microscopic interactions corresponding to two distinct binding modes, namely upper-stem and lower-stem bound conformations respectively.

Beyond barrier height, the free energy profiles illuminate distinct landscape features. HIV-2 displays a broad single minimum centred at a mean COM distance, while HIV-1 reveals a broader global well subdivided into two basins separated by a shallow barrier. Ensemble structures extracted from unbiased simulations (**Figure 2C–E**) further resolve these differences: HIV-2 maintains a dominant binding mode, whereas HIV-1 supports two coexisting conformations-an upper-stem bound global minimum and a lower-stem metastable state. Consistent with this, equilibrium simulations of HIV-1 yield a bimodal COM distance distribution (**Figure S7**), whereas HIV-2 fluctuates around a single mean (∼0.6 nm). Thus, the C24-induced bulge expansion in HIV-1 not only elevates the dissociation barrier but also introduces an accessible intermediate state under ambient conditions.

To dissect the microscopic determinants of these divergent landscapes, we analysed residue-level contact probability maps **(Figure S4)**. Despite structural similarity, the two TAR–Tat complexes adopt distinct binding postures. In HIV-2, the NMR structure (PDB id: 6MCE(38)) reveals a bulge-centred sandwich formed by Arg49, U23, and Arg52. During equilibration, however, the Arg52-U23 contact weakens, leaving only Arg49-U23 intact but reoriented, which drives Tat toward a lower-stem inclined position with minimal loop engagement (**Figure 2C**, **zoom**). In contrast, HIV-1 Tat alternates between two modes: a coiled lower-stem conformation (**Figure 2E**) and an extended upper-stem bound conformation (**Figure 2D**). In the coiled state, Arg52 sustains its interaction with U23, while Lys50 dynamically engages loop residue G34, but Arg49 fails to stabilize bulge contacts, limiting loop connectivity (**Figure 2E, zoom**). In the extended conformation, however, C24 insertion restores Arg49-U23 contact, forming a pseudo-sandwich with Arg52. This rearrangement lifts Tat upward, enabling cooperative bulge-loop engagement (**Figure 2D, zoom**). Together, these findings show that a single bulge insertion fine-tunes local binding topology, reshapes Tat binding modalities, and alters distal site accessibility, thereby diversifying RNA-protein recognition pathways.

### Active Site Bulge Mutation Alters Protein Binding Mechanism: Switching from Conformational Selection to Induced Fit

So far, our analysis has mapped the distinct molecular interactions across TAR variants, yet the fundamental binding mechanism remains unresolved. To uncover this, it is essential to consider the dynamics of the apo RNA, which serves as a structural ‘propeller’ preorganizing conformational ensemble that dictates distinct binding modes. In HIV-2 apo TAR, the loop and bulge engage in strong dynamic communication **(Figure 1)**, reinforced by U23-A35 base stacking (**Figure S8**). The centre-of-mass distance between these two domains exhibits a bimodal distribution **(Figure 3A)**, corresponding to a stacking-induced proximal state **(Figure 3B)** and an A35-extrusion-induced separated state **(Figure 3C)**. Our analysis reveals that the presence of a distorted U23–A27:U38 base triple **(zoom Figure 3B-C)**, representing a preorganized conformation within the HIV-2 apo ensemble, is primarily contingent upon this dynamic U23-A35 interaction. This stacking-driven breathing motion narrows and widens the RNA major groove, creating a dynamic conformational selection mechanism(36, 60, 61) for Tat recognition. Under this regime, Tat is preferentially accommodated in lower-stem inclined conformation, with minimal loop engagement (**Figure 3D**). By contrast, in HIV-1 TAR, the larger and more heterogeneous bulge(32) (**Figure S8**) abolishes the U23-A35 stacking interaction. This enforces a persistently greater spatial separation between loop and bulge (**Figure 3E)**, yielding an inherently wider major groove (**Figure 3F**), thereby reducing the propensity for base triple formation in the apo ensemble **(zoom Figure 3F)**. Weeks and Crothers previously proposed that Tat accommodation in HIV-1 TAR occurs via groove widening(24, 32, 62) enabled by bulge plasticity.

**Figure 3:**
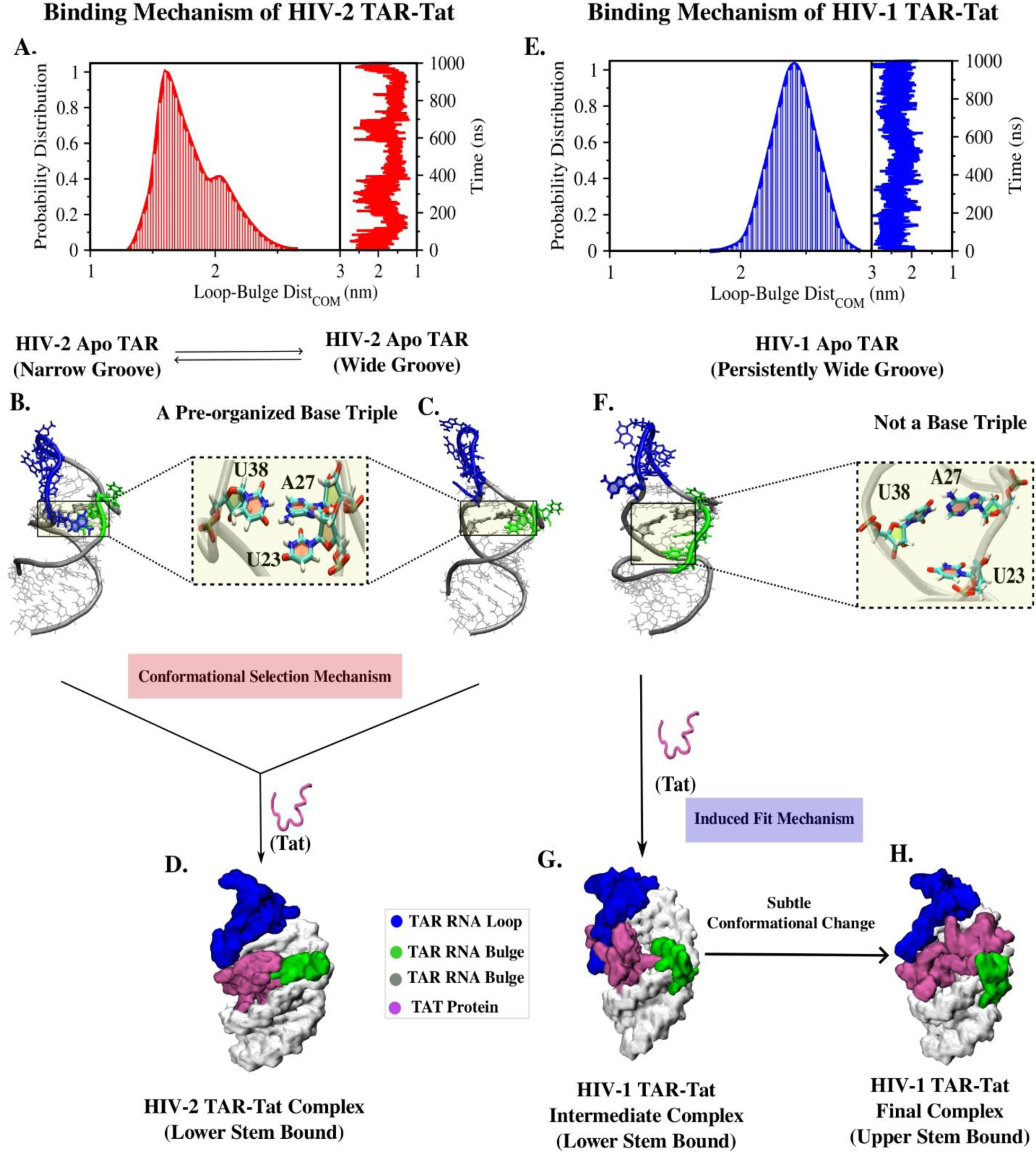
Microscopic binding mechanism. **(A)** represents the centre of mass distance between loop and bulge along the trajectories and its probability distribution in HIV-2 apo TAR RNA. **(B)** and **(C)** illustrate a breathing between A35-U23 base-stacked and unstacked conformations, dynamically preorganizing specific base triple formation U23–A27:U38 **(D)** is a MD snapshot representing a lower stem inclined HIV-2 TAR-Tat complex. **(E)** represents the centre of mass distance between loop and bulge along the trajectories and its probability distribution in HIV-1 apo TAR RNA. **(F)** shows a large and heterogeneous bulge discarding base-triple formation. **(G)** is a MD snapshot for an intermediate lower stem inclined conformation in HIV-1 apo TAR RNA. **(H)** is a snap out of MD illustrating the upper-stem bound stable HIV-1 TAR-Tat binding mode, representing an extended conformation of Tat, interacting to both bulge and loop simultaneously.

Consistent with this view, the widened groove provides Tat an enhanced translational mobility along the RNA helical axis, allowing it to explore binding sites until initial stabilization occurs via a U23-Arg52 tether. This interaction anchors a lower-stem bound intermediate conformation (**Figure 3G**), which then promotes conformational rearrangements at the primary bulge-binding site. Specifically, Tat binding facilitates formation of the base triple(21, 22) (U23–A27:U38)-a conformational feature absent in the apo state. As binding interactions mature, Arg49 joins U23 and Arg52 to form the pseudo-sandwich motif (**Figure 2**), subtly repositioning Tat upward toward the upper stem and enabling loop engagement (**Figure 3H**). This successive progression exemplifies an induced fit mechanism(63). Tat initially stabilizes a weaker lower-stem conformation, then actively stimulates RNA rearrangements that culminate in a highly stabilized upper-stem bound complex.

### Post-Tat-Binding Allosteric Interplay between Loop and Bulge Promotes Cascade Protein Binding Event

In HIV-2 TAR, Tat binding is geometrically restricted to the lower stem and primarily interact with A35, with minimal engagement of the loop (**Figure 4A**). By contrast, HIV-1 TAR supports additional loop interactions, most notably with A35 and G34, including the experimentally validated Lys50–G34 contact(64) (**Figure 4B**). The absence of stable loop interactions in HIV-2 destabilizes loop dynamics, generating a distinct fluctuation pattern. In our earlier work, we proposed that this reconfigured loop in HIV-2 TAR, upon Tat binding, activates a new fluctuation centred on residue G32 (**Figure S9**). When flipped outward, this residue likely becomes a binding-competent site for TAR’s secondary partner, Cyc-T1. Thus, in HIV-2, the loop and bulge function as dynamically coupled motifs, allosterically primed to promote subsequent protein binding events. In striking contrast, Tat binding to HIV-1 TAR directly stabilizes the loop. Here, the strong Tat–loop interactions, integrating A35 and G34, promote the flipped-out conformation of G32, whereas the flipped-in state is likely restricted to avoid steric crowding within the loop as surmised from the base-stacking probability map as drawn following Barnaba definition (**Figure S10**). This outward projection of G32 is likely to establish a conformationally favorable capture mode for downstream binding of another protein, Cyc-T1. Pseudo-dihedral angle measurements (**Figure 4C**) reveal a bimodal distribution of G32 base-flipping events in HIV-2 TAR, reflecting a dynamic equilibrium, whereas in HIV-1 TAR–Tat complexes the unimodal distribution indicates that G32 is uniquely stabilized in an outward-exposed state (**Figure 4D**).

**Figure 4:**
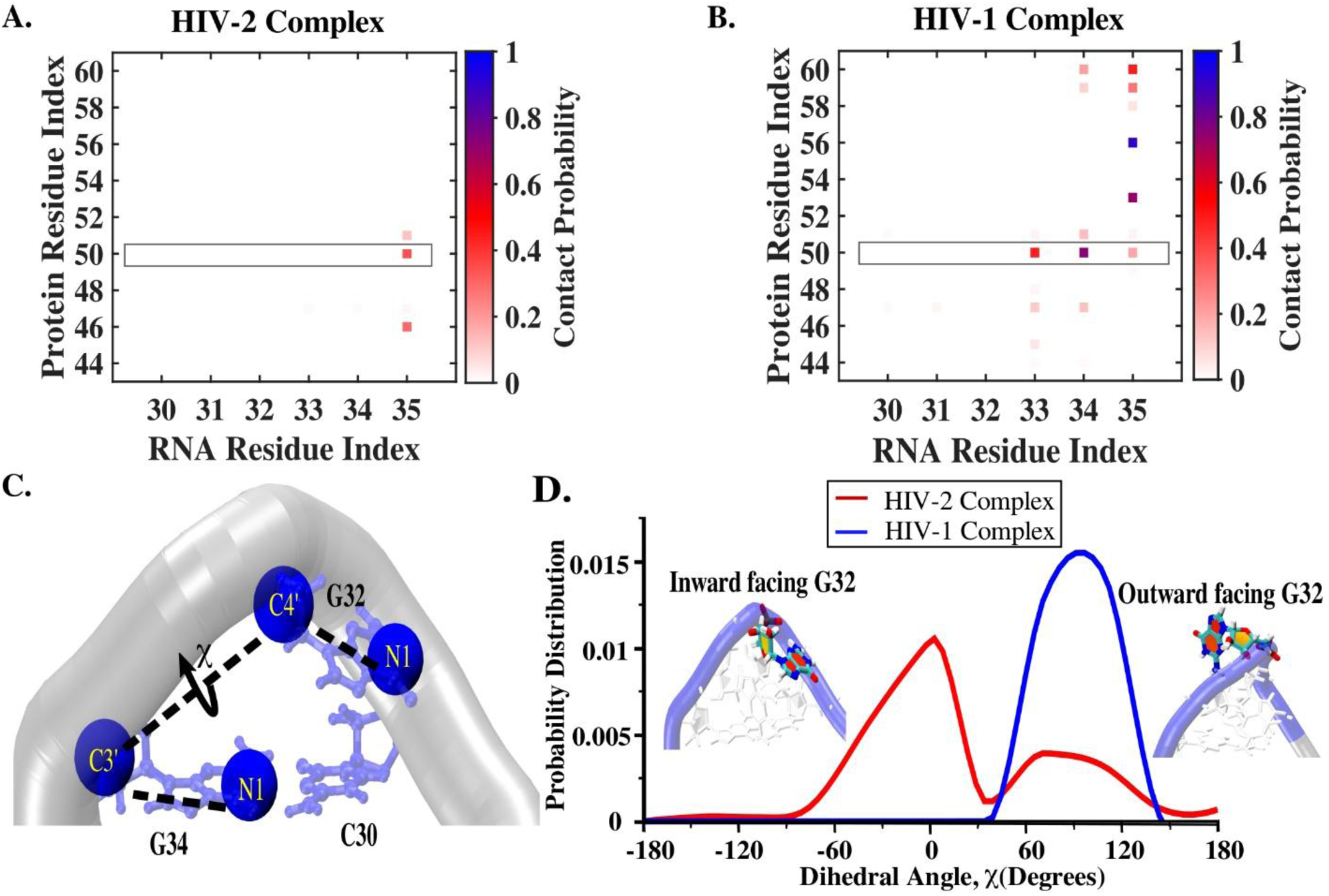
Comparison of post-Tat-binding Allosteric Interplay between HIV-2 TAR and mutant TAR (HIV-1). **(A) and (B)** represent the contact probability map for loop and Tat interaction drawn at residue level in HIV-2, HIV-1 TAR-Tat respectively. The black rectangular box highlights the contact pair developed by Lys50 (Tat) and loop residues of TAR RNA. **(C)** represents a pseudo-dihedral angle (χ), schematically drawn to define the flip in/out dynamics of G32**. (D)** compares probability distribution of pseudo-dihedral angle (χ) between HIV-2 vs HIV-1 in their respective complex forms.

### Navigating Shortest Allosteric Pathways from Bulge to Loop and Pathway-Switching Upon Single-Point Mutation and Protein Binding

Taken together, our findings suggest that in HIV-2, the more dynamic binding of Tat to TAR RNA reorganizes local fluctuations and facilitates recruitment of the next cascade protein, Cyc-T1, through a signal transduction mechanism that searches for the optimal conformational capture required for next protein’s recognition. In contrast, in HIV-1, the more integrated Tat–TAR binding promotes subsequent protein engagement by activating optimal conformation, likely to facilitate more stable binding. Regardless of the viral variant, transcription necessitates Cyc-T1 binding near G32 following Tat association at the bulge, highlighting an intrinsic loop-bulge allosteric coupling that functions as a conduit for signal transduction. Here, the bulge and loop act as remote information hubs (sources and sinks).

To essentially understand how a single-point mutation (C24 insertion in HIV-1) reshapes this communication, we applied a graph-network based algorithm inspired by Dijkstra’s shortest-path approach. This method combines position-based fluctuation correlations with heavy-atom contact probabilities, assigning nodes and edges to trace information flow from the bulge (source) to the loop (sink), thus identifying the shortest pathways of communication. In HIV-2 apo TAR, two distinct pathways emerge: (i) a canonical route mediated by classical local structure-based interaction along a single strand of the stem, and (ii) a noncanonical pathway dynamic non-local interaction between U23 and A35 (**Figure 5A, 5B**), consistent with our contact PCA analysis. In HIV-1 apo TAR, the C24-induced bulge heterogeneity suppresses the U23–A35 dynamic stack and significantly perturbs global dynamics (**Figure S5**), redirecting signal propagation through zigzag canonical local contacts spanning both strands of the stem (**Figure 5C**). Upon Tat binding, communication pathways are rewired in both variants. In HIV-2, Tat disrupts U23–A35 stacking, eliminating the noncanonical route and leaving only the canonical pathway intact (**Figure 5D**). In HIV-1, Tat binding dampens overall TAR dynamics, constraining information flow primarily along a single-strand pathway (**Figure 5E**).

**Figure 5:**
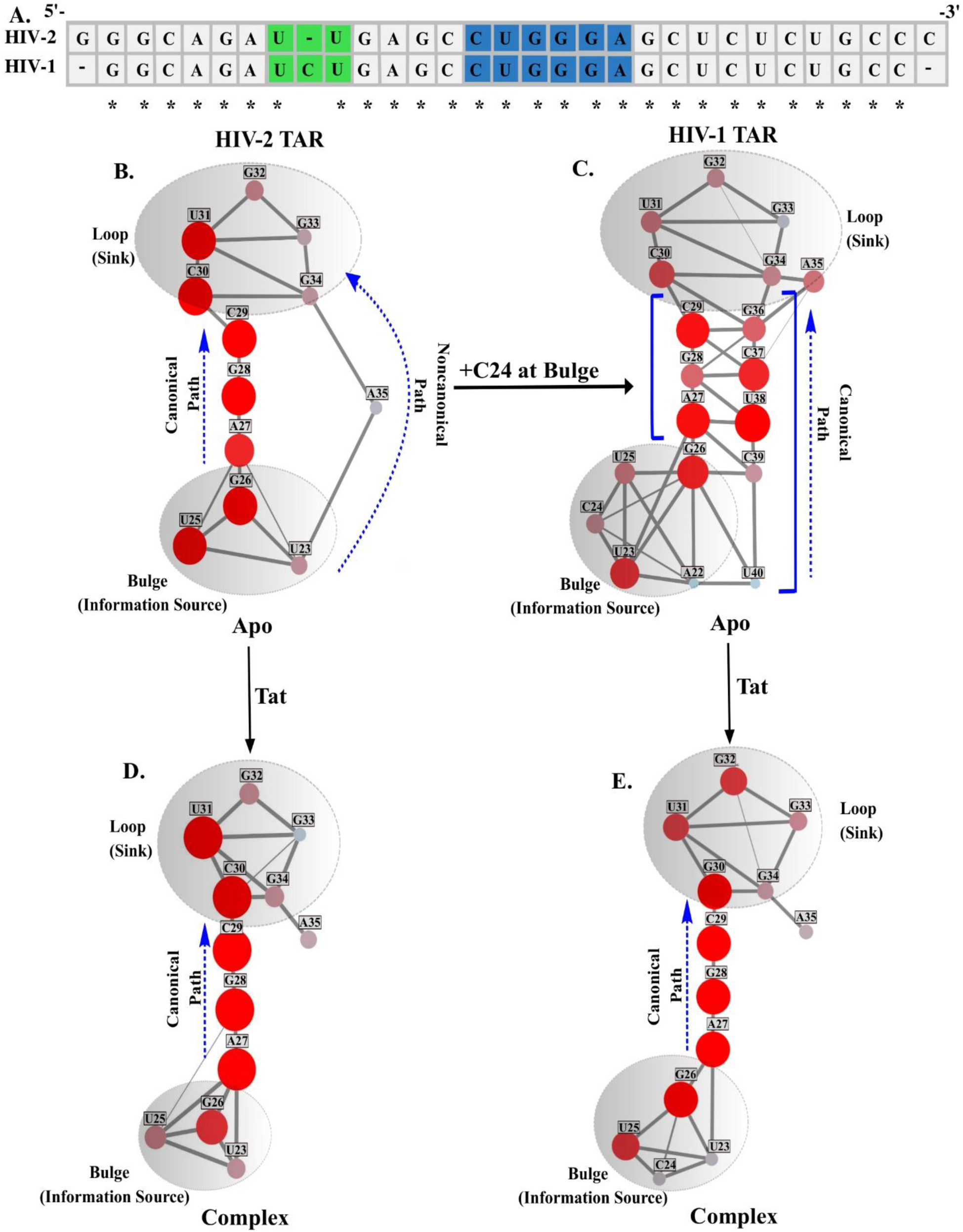
Effect of a single point mutation on functional mode of allosteric interplay between loop and bulge. **(A)** depicts sequence alignment between TAR RNAs of HIV variants, reflecting that HIV-1 and HIV-2 are separated by a single point (C24 at bulge) mutation. **(B–E)** Allosteric signalling information channels, derived employing graph-network-based shortest path algorithm, between bulge (source) and loop (sink) as shown by grey coloured hubs in HIV-2 apo TAR (B), HIV-1 apo TAR (C), HIV-2 complex TAR (D), and HIV-1 complex TAR (E). Each node here presents the nucleotide index, while the edges connecting any two nodes are weighted by the frequency of highly correlated nucleotide pairs. The size/brightness of nodes indicates their recurrence across the dominant communication paths between source and sink. The dotted blue arrows present the flow of information from source (bulge) to sink (loop) in each case.

## CONCLUSION

Since 1998(13), in vitro selection strategies such as SELEX have generated TAR- and Tat-binding aptamers, including RNA^Tat and related DNA/RNA variants that inhibit viral replication. While these studies underscore the therapeutic relevance of the TAR–Tat interface, our work focuses on the mechanistic principles by which RNA sequence and architecture encode recognition, leveraging a systematic, computational framework. A central theme emerging from our study is that fluctuating RNA motifs, particularly bulges and loops, act as intrinsic regulators of binding. By shaping structural flexibility and governing interdomain communication, these motifs channel Tat into distinct recognition pathways. Importantly, even single-nucleotide changes can remodel this regulatory network, illustrating how RNA encodes molecular adaptability at the most granular scale.

The methodology, combining tens of microseconds of atomistic and free-energy simulations at physiological salt environment, with contact-based PCA and graph-network analysis, enables the first systematic mapping of allosteric communication pathways and their rewiring upon mutation or protein binding. Comparative analyses across HIV-1, HIV-2, and BIV TAR reveal four key mechanistic insights:

(i) First, bulges and loops function as regulatory modules rather than passive irregularities. In HIV-2, strong U23–A35 stacking and loop–bulge communication constrains Tat to a lower-stem posture, whereas the larger HIV-1 bulge disrupts this preorganization, widens the major groove, and enables loop participation. Such motif-driven tuning positions local flexibility as the “design logic” of TAR recognition.
(ii) Second, a single nucleotide can rewire the entire recognition landscape. The C24 insertion in HIV-1 alters bulge–loop communication, destabilizes base triples, and generates new cooperative contacts between Tat residues Arg49 and Arg52. This change allows Tat to transition from a metastable, lower-stem encounter to a higher-affinity upper-stem binding mode. Thus, minimal sequence edits can fine-tune recognition by reshaping RNA dynamics.
(iii) Third, different TAR variants embody distinct binding mechanisms. HIV-2 relies on conformational selection(36), where Tat exploits preorganized structural states. HIV-1, in contrast, follows an induced fit pathway, with Tat actively remodelling the RNA upon binding. The induced fit mechanism(63) is consistent with the finding of several early works (65–67). BIV TAR, with reduced self-integrity, achieves high affinity to Tat. These divergent strategies underscore how evolutionary changes in RNA sequence dictate fundamentally different energetic and kinetic routes to recognition.
(iv) Finally, loop-bulge communication emerges as a powerful allosteric control element. In HIV-2, Tat binding destabilizes the loop, creating new fluctuation modes at G32 that likely prime Cyc-T1 recruitment. In HIV-1, Tat instead stabilizes the loop and outwardly projects G32, generating a conformational state directly competent for downstream protein capture. These results show how local fluctuations propagate into distal effects, embedding regulatory logic within RNA architecture.

Together, these findings not only highlight sequence-encoded allosteric control as a determinant of RNA-protein recognition, but also provide a systematic mapping of RNA allostery. By identifying key regulatory motifs and pathways, this approach complements experimental strategies like SELEX, enabling focused sequence-space exploration and rational design of high-affinity RNA–protein interactions.

Beyond HIV, these principles reveal that viral and regulatory RNAs generally encode modular allosteric “control knobs,” where subtle sequence changes can reprogram recognition and function. Harnessing such design rules offers a predictive framework for RNA engineering and targeted therapeutic development, moving from descriptive to mechanistic understanding.

## Supporting information

https://drive.google.com/file/d/1spYDiWt2stml7pFnI3tTT5L_TjtT033Y/view?usp=sharing

## AUTHOR CONTRIBUTION

DS and AC contributed equally to this paper. Conceptualization: SR; data curation: DS and AC.; formal analysis: AC, DS, AS, RS, SR; funding acquisition: SR; methodology: AC, DS, AS, RS, SR; project administration: SR; resources: SR; validation: AC, DS, AS, RS, SR; writing-original draft: DS, AC, SR; writing, review and editing: AC, DS, AS, RS, and SR.

## DECLARATION OF INTERESTS

The authors declare no competing interests.

## ACKNOWLEDGEMENTS

SR acknowledges support from the Department of Biotechnology (DBT) (Grant No. BT/12/IYBA/2019/12 and BT/PR40192/BTIS/137/69/2023). DS acknowledges UGC fellowship and AC and AS acknowledge DST INSPIRE fellowship.

## Notes

### Competing Interest Statement

The authors have declared no competing interest.

