## Supplementary material for "Mapping Allosteric Rewiring in Viral RNA: Sequence-Encoded Control of Protein Binding Mechanisms": https://drive.google.com/file/d/1spYDiWt2stml7pFnI3tTT5L_TjtT033Y/view?usp=sharing

### This PDF file includes:

**Supporting Information**

- **Supporting Method**
  - SM1: System preparation for BIV
- **Supporting Figures**
  - Figure S1 to S10
- **Supporting Table**
  - Table S1 to S6
- **References for Supporting Information**

### Supporting Method

#### SM1: System Preparation for BIV

The initial coordinates of the TAR-Tat complexes for BIV (1BIV (1)) were directly obtained from the Protein Data Bank, representing the solution NMR structures reported by Patel et al.. For the BIV complex, among the five available NMR models, model-1 was selected since all the model structures exhibit close structural resemblance. The corresponding free TAR RNA structures were derived by selectively removing the Tat peptide coordinates from their respective RNA grooves.

### Supporting Figures

#### Sequence, structure and structural integrity: Comparing TAR-Tat complexes of three different variants:

Interaction energy is a macroscopic quantity that reflects the structural integrity of a system and the degree of stabilization between two molecular entities, governed by the underlying interaction potential at either the intra- or intermolecular level. To evaluate specific interaction energies between RNA-RNA and RNA-protein pairs, the gmx energy module in the GROMACS(2) package is generally used. By selecting the relevant energy terms, including short-range (SR) and 1-4 electrostatic (Coulombic) as well as Lennard-Jones (LJ) interactions, we can extract the time-resolved interaction energy profiles for the chosen molecular pairs. The total interaction energy is obtained by summing the corresponding Coulombic and LJ components over the course of the simulation. This energy is recorded in the edr file, which stores all energy components calculated based on the parameters defined in the mdp file and the employed force field.

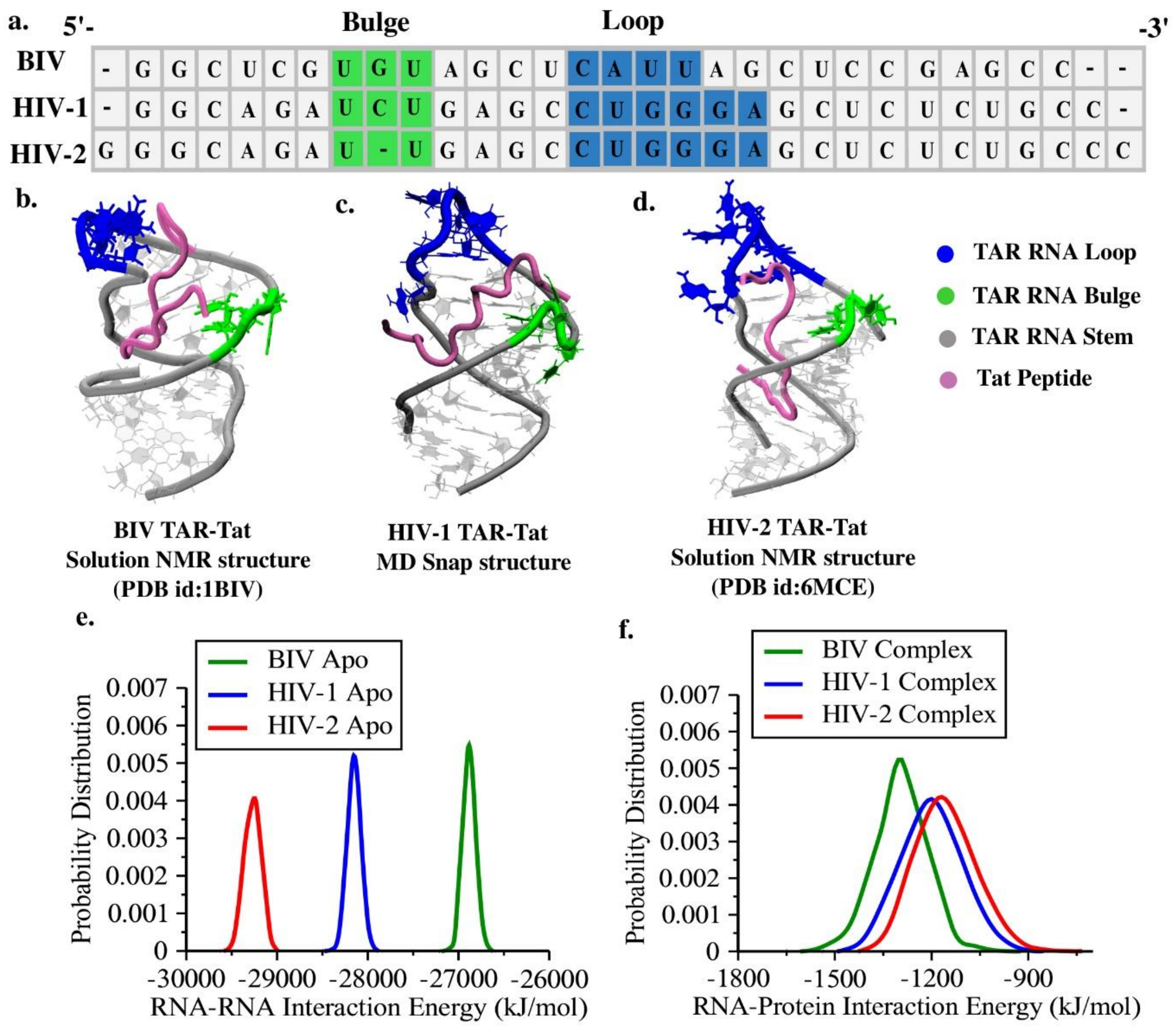


**Figure S1**: **Comparison of the intra-molecular interactions in RNA and inter-molecular interactions between RNA and protein among variants**: **(a)** depicts the primary sequences in TAR RNA variants BIV, HIV-1 and HIV-2 respectively. **(b)**, **(c)** and **(d)** show respective TAR-Tat complex structures. **(e)** represents the probability distribution of intra-molecular internal energy within BIV, HIV-1 and HIV-2 RNAs. **(f)** depicts the probability distribution of inter-molecular interaction energies between TAR RNA and Tat protein for BIV, HIV-1 and HIV-2.

**Comparing spatial packing of surrounding water and ions around TAR RNAs in apo and complex states across the variants:**

Determining the correct ionic concentration requires locating the bulk region of ions and water along the radial shells surrounding the RNA.

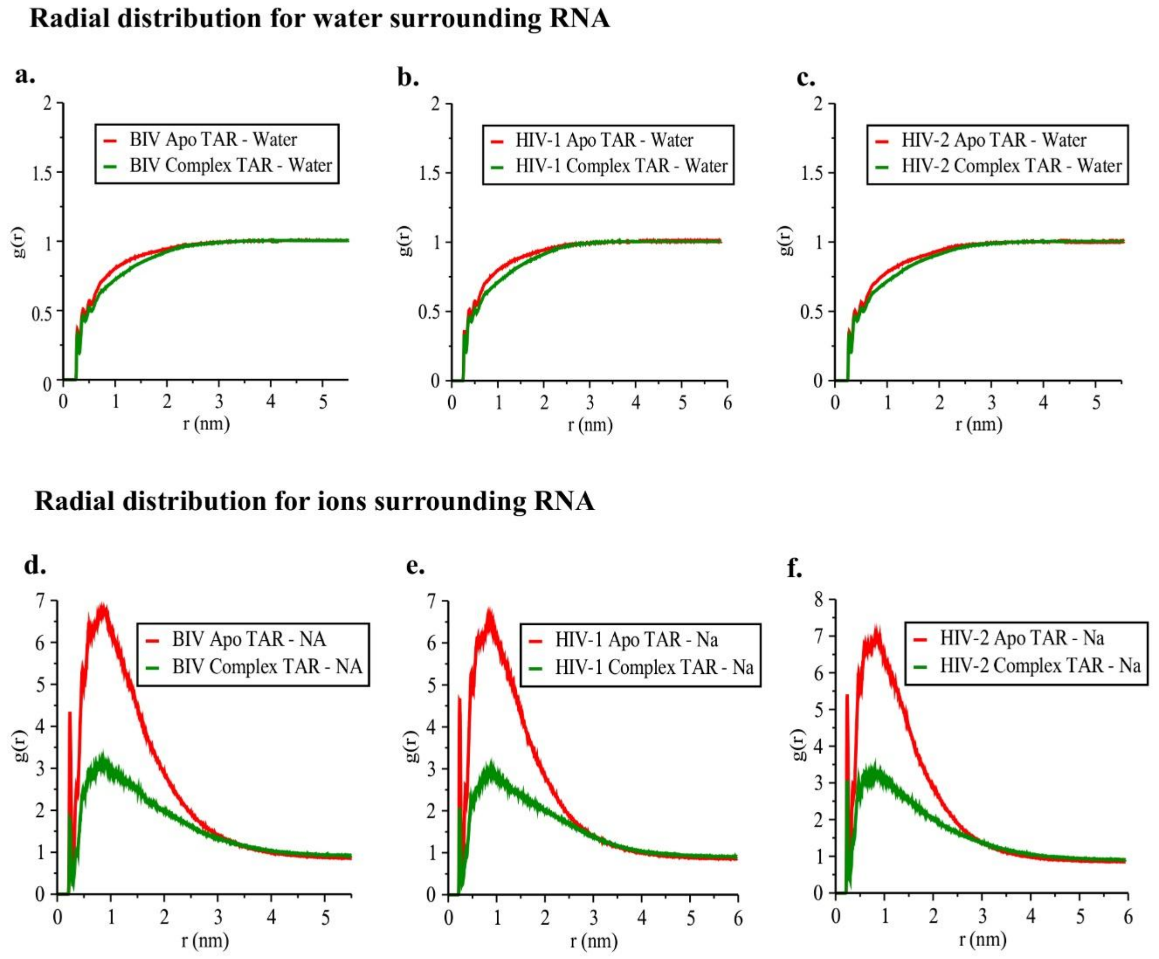


**Figure S2: Radial distribution function (RDF) plots: (a), (b) and (c)** represent the RDF profiles of water molecules around TAR RNA in BIV, HIV-1 and HIV-2 respectively**. (d), (e) and (f)** represent the RDF profiles of $Na^{+}$ ions around TAR RNA in BIV, HIV-1 and HIV-2 respectively. Only heavy atoms are taken into consideration.

**Comparison of free-energy profile across the variants along with respective windows overlapping for HIV-2 and HIV-1 TAR-Tat:**

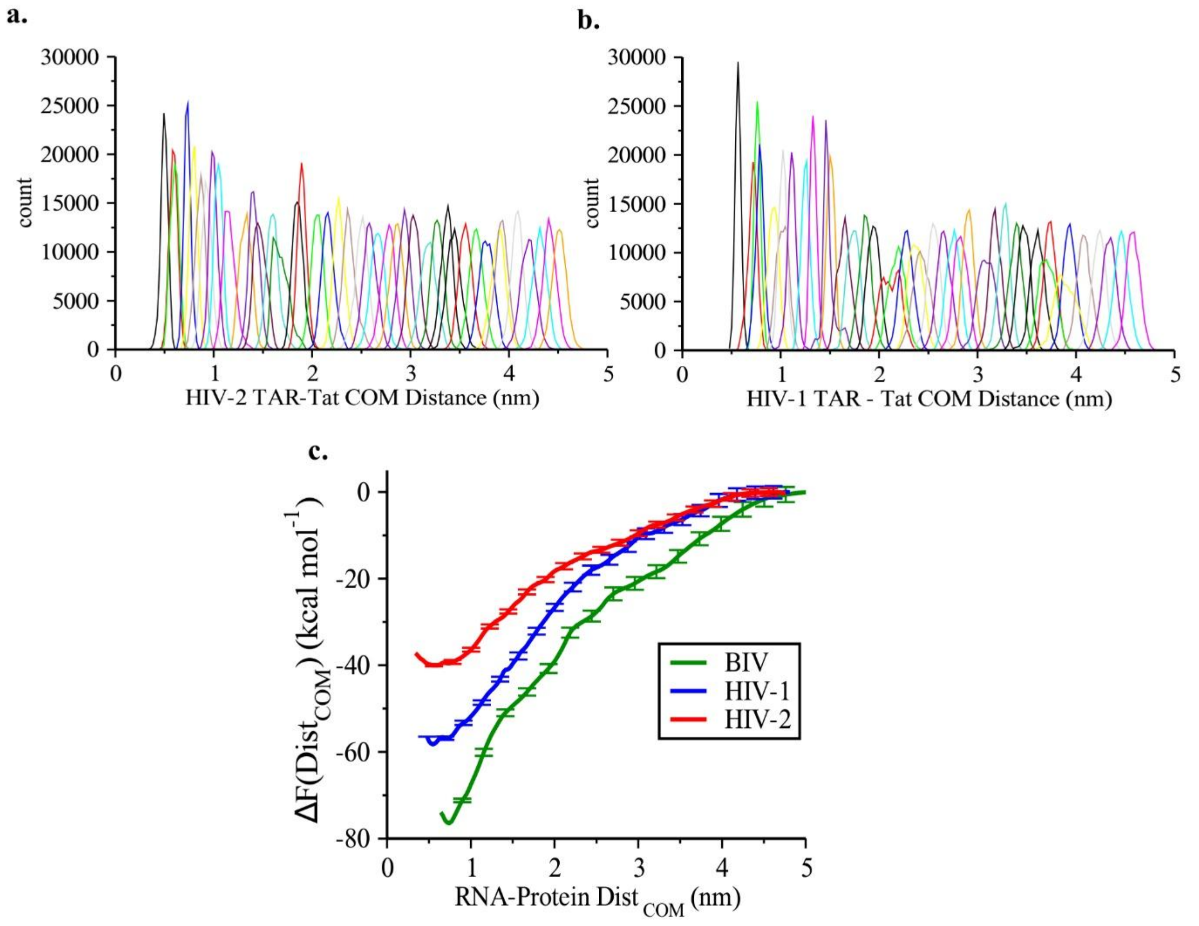


**Figure S3: Comparison of free-energy profile along with respective windows overlapping for HIV-2 and HIV-1 TAR-Tat dissociation: (a)** shows 42 windows overlapped well along the COM distance of HIV-2 TAR and Tat, **(b)** shows 42 overlapping windows covering the whole sampling phase space along the COM distance of HIV-1 TAR and Tat and **(c)** represents the comparison of free energy of TAR-Tat dissociation across different variants.

**Radial distribution function for all the heavy atoms of RNA w.r.to all the heavy atoms of protein and all atom inter-chain contact probability map**

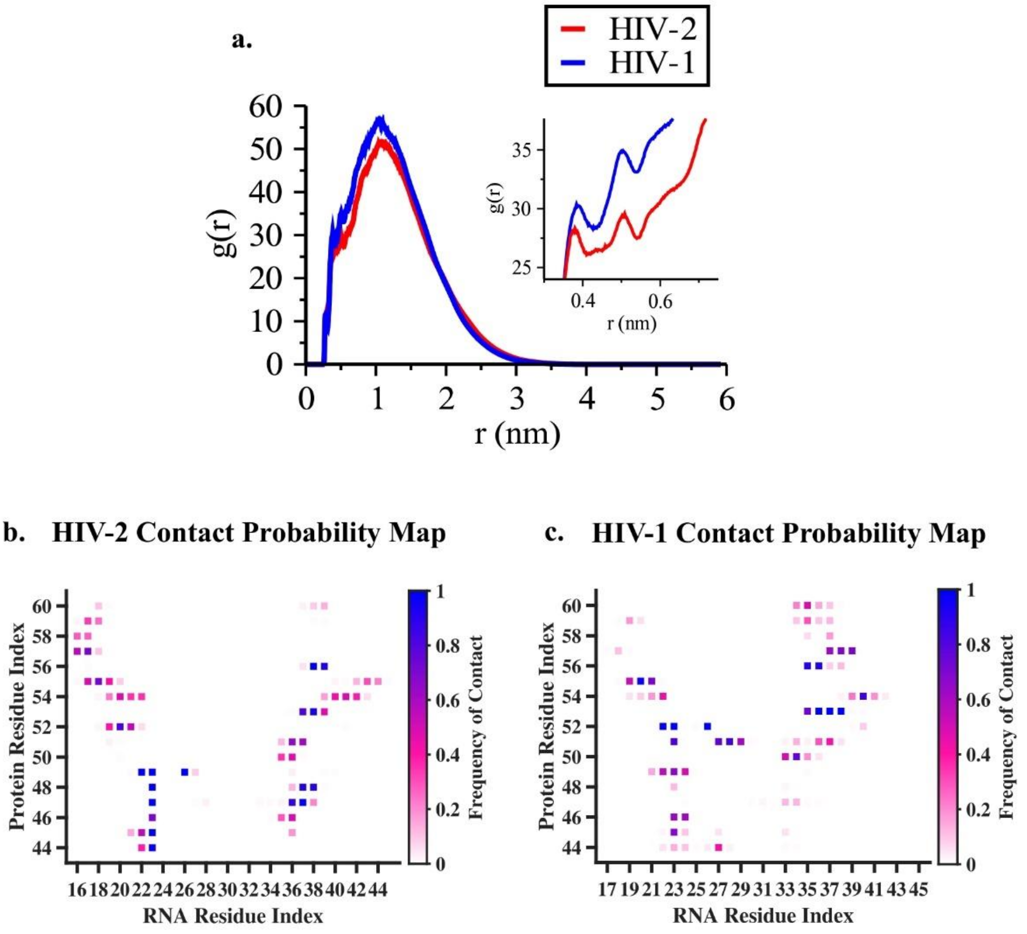


**Figure S4: Analyses of RNA-protein spatial interactions: (a)** represents the Radial distribution function (RDF) of RNA around protein's heavy atoms, used to define the contact threshold. **(b) and (c)** represent residue wise contact maps between RNA and protein in HIV-2 and HIV-1 respectively, based on probability of contact formation along the trajectories.

**Positional Fluctuation based Covariance Analysis comparing apo and complex states (at all heavy atoms level) across the variants:**

To quantify correlated motions across the RNA, we construct the covariance matrix of atomic positional fluctuations. For any two heavy atoms *a* and *b*, the covariance is defined as

$$C_{ab} = \langle \Delta \mathbf{r_a} . \Delta \mathbf{r_b} \rangle \qquad eq. 1$$

$\Delta \mathbf{r_a}$ and $\Delta \mathbf{r_b}$ are the displacement vectors of atoms *a* and *b* from their mean positions, and $\langle \cdot \rangle$ denotes the time average over the trajectory. The diagonal elements (a=b) capture the mean-square fluctuations of individual atoms, while the off-diagonal elements (a≠b) report on the degree of correlated or anticorrelated motion between atomic pairs. The range of covariance values have been shown in $nm^2$ within the range of -0.2 to 0.2 $nm^2$ we compared our four sets.

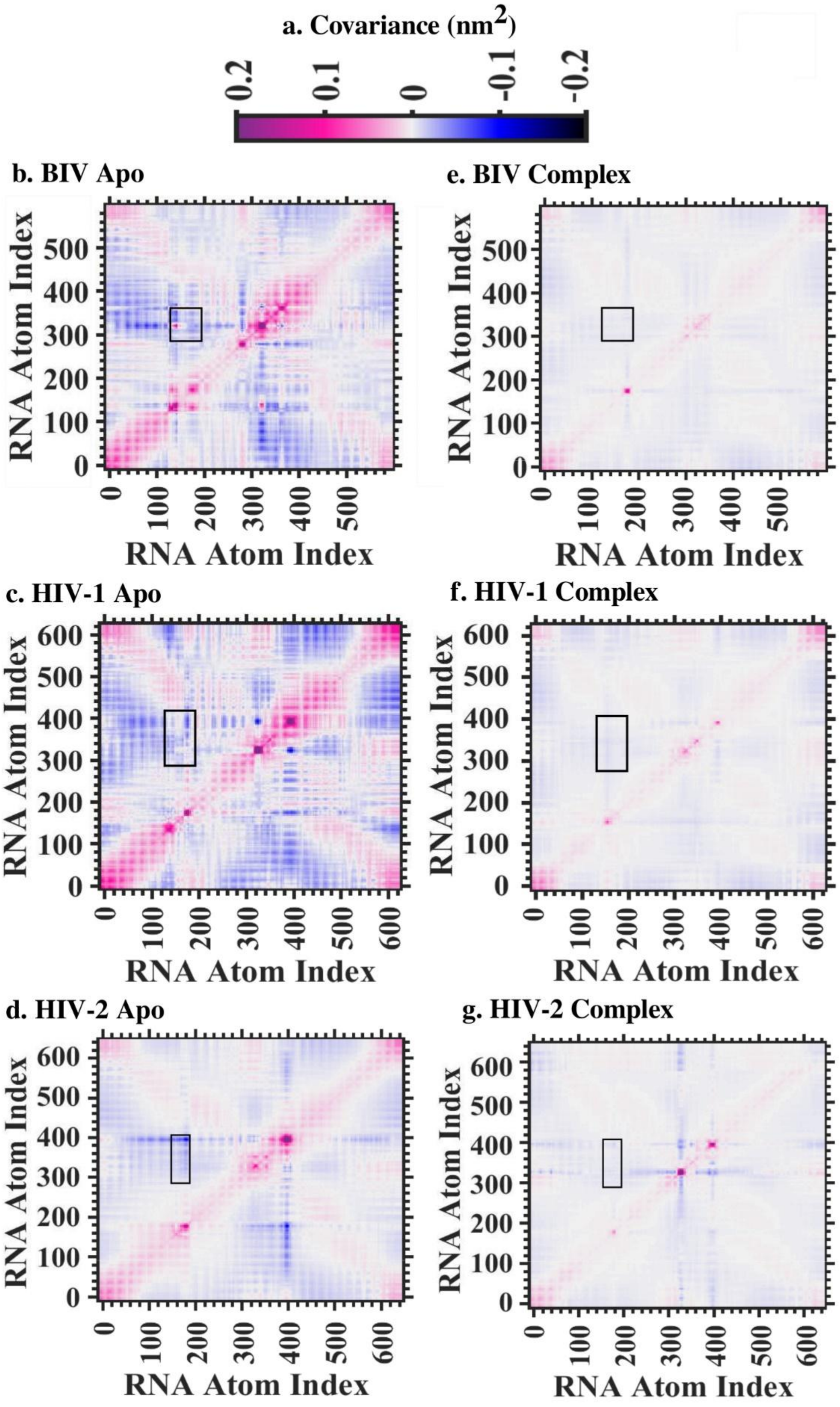

**Figure S5: Covariance matrix for all three variants' TAR RNAs for both their apo and complex form:** **(a)** represents the colour bar standing for covariance within the range +0.2 to -0.2. **(b), (c) and (d)** are the respective covariance matrices for apo TAR RNAs of BIV, HIV-1 and HIV-2. **(e), (f) and (g)** are the similar matrices in the corresponding Tat bound states for the BIV, HIV-1 and HIV-2 respectively.

**Root Mean Square Deviation (RMSD):**

RMSD is an essential parameter to quantify the extent to which a molecular segment deviates from its reference or native conformation over time. It provides insight into whether the molecule has transitioned into a distinct conformational state during the simulation. RMSD is computed by averaging the squared deviations of atomic coordinates from the reference structure, offering a time-resolved measure of structural changes. For biomolecular system, the calculation is typically mass-weighted to account for the contribution of each atom. The RMSD at a given time t is expressed as:

$$RMSD(t) = \sqrt{\frac{1}{M}\sum_{i=1}^{N} \mathrm{m_i}|x_i(t) - x_i(0)|^2} \qquad eq.2$$

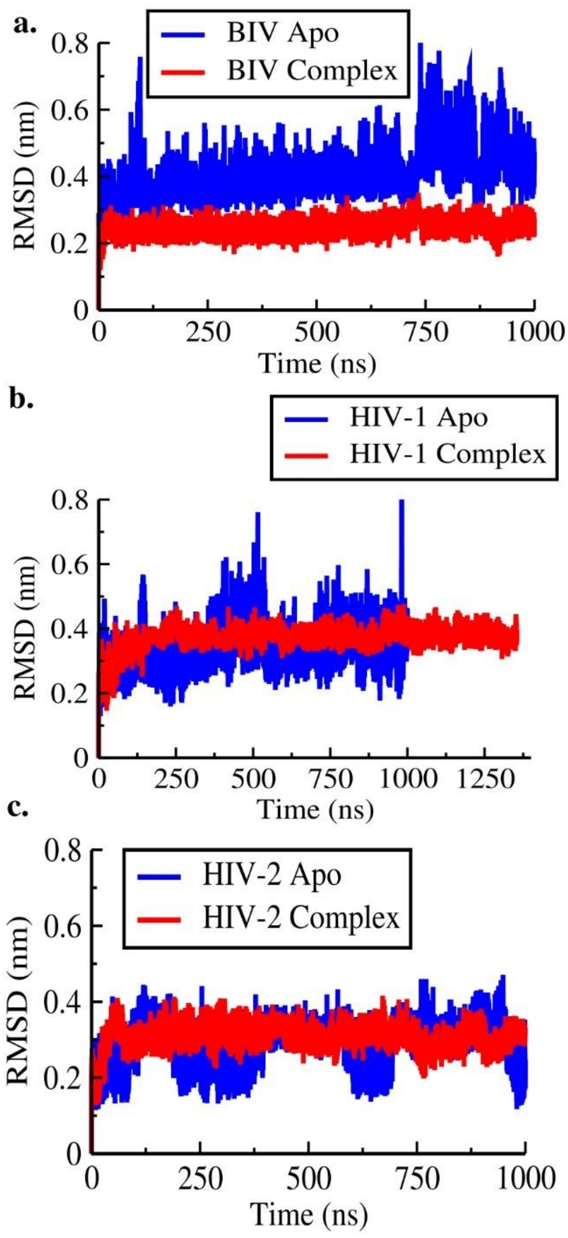


**Figure S6: Comparison of RMSD along the trajectories: (a), (b) and (c)** represent the RMSD of TAR RNA in apo vs complex state for BIV, HIV-1 and HIV-2 respectively.

**Probability Distribution of center of mass (COM) distance btween TAR and Tat in HIV-2 vs. HIV-1:**

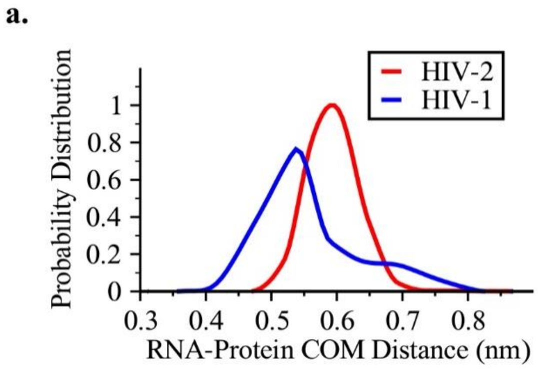


**Figure S7: Comparing the RNA-protein COM distances:** This panel represents the COM distances between RNA and protein along the trajectories for HIV-2 and HIV-1 complex.

**Comparing loop-bulge communication of HIV-2 to HIV-1 apo TAR:**

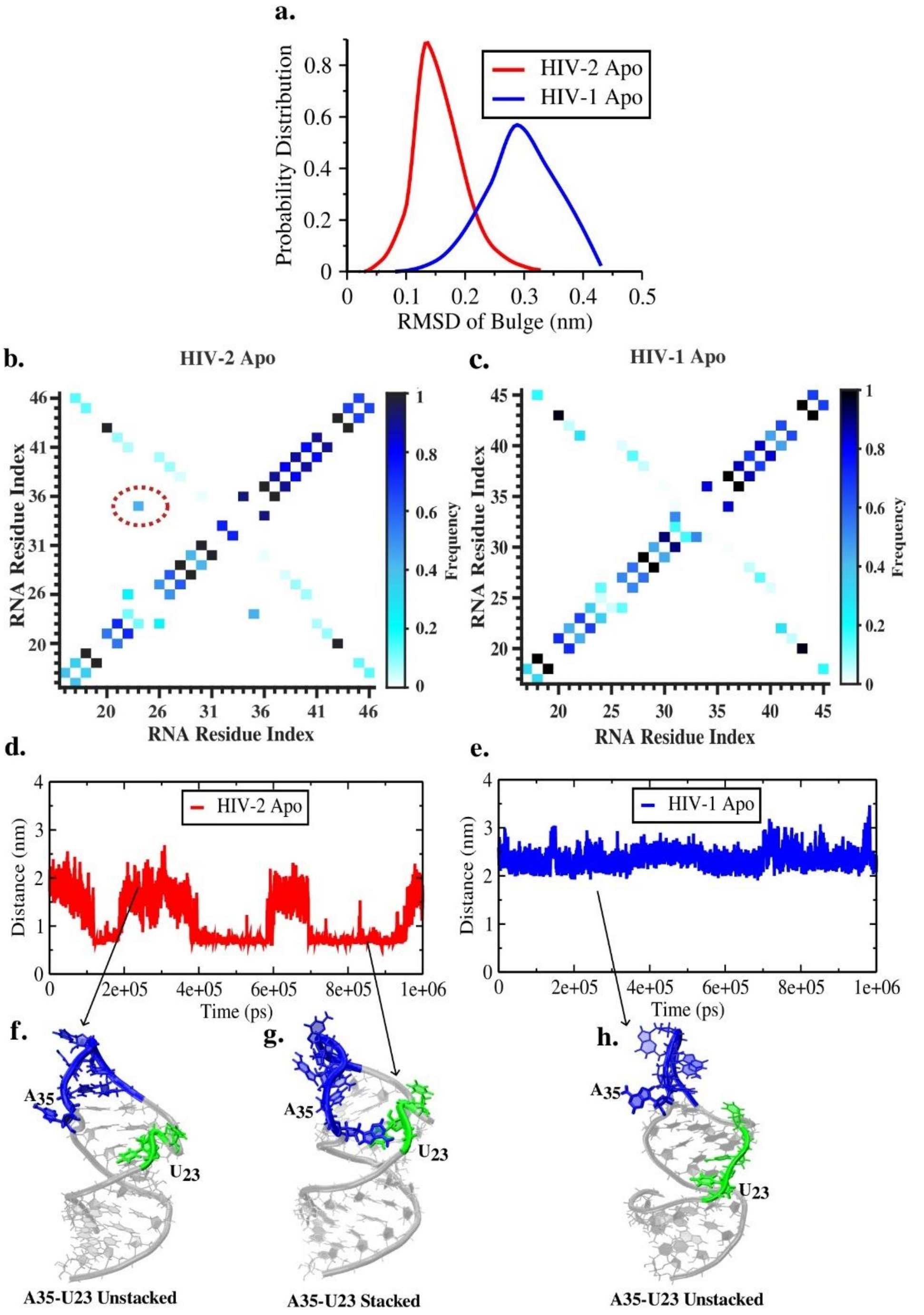


**Figure S8: Distal loop-bulge communication in apo TAR RNAs: (a)** represent the RMSD of bulge interpreting bulge dynamicity **(b) and (c)** illustrate the RNA base stacking interactions with corresponding frequency of occurrence calculated by Barnaba python library(3) observed in apo HIV-2 and HIV-1 respectively. A critical U23(from bulge)-A35(from loop) base stacking interaction mediating loop-bulge communication in HIV-2 TAR is highlighted with dotted circle. **(d) and (e)** represent their (U23 and A35) centre of mass distance along the trajectories of HIV-2 and HIV-1 apo TAR RNA respectively. **(f) and (g)** are the representative structures of HIV-2 apo TAR while A35 and U23 are unstacked and stacked respectively. **(h)** is the only possible unstacked conformation of HIV-1 apo TAR with respect to the loop-bulge (A35-U23) communication.

**Comparative RMSF for apo to complex state of TAR RNAs:**

RMSF provides a quantitative measure of the temporal fluctuations of individual residues or atoms within a protein throughout the simulation. It is particularly effective in distinguishing between flexible (unstructured) and rigid (structured) regions of a biomolecule. RMSF offers residue-specific insights by averaging positional deviations over time. The fluctuation for atom i is given by:

$$RMSF_i = \sqrt{\frac{1}{T} \sum_{\tau=1}^{T} |x_i(\tau) - \bar{x}_i|^2} \qquad eq.3$$

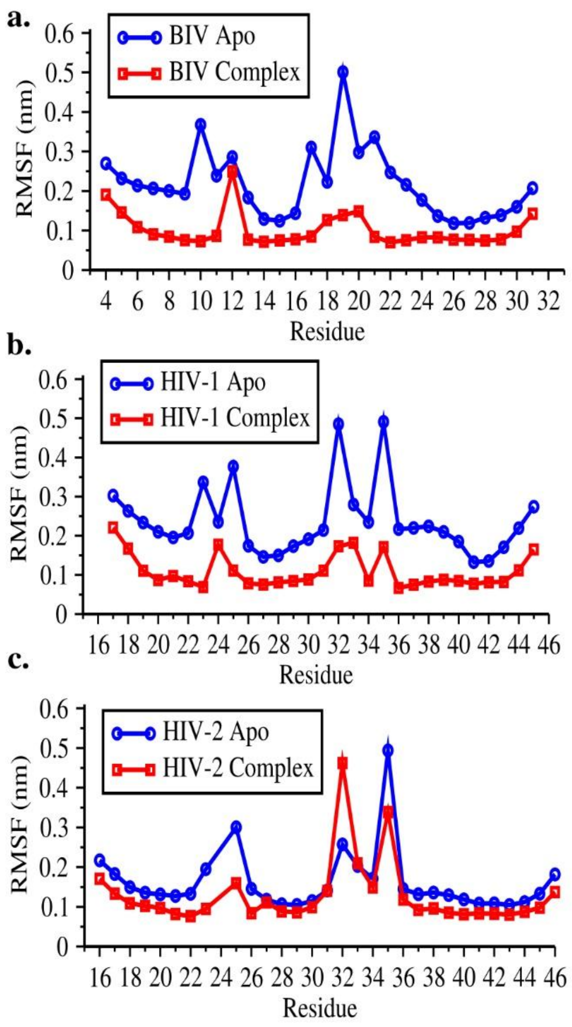


**Figure S9: Comparison of RMSF Plots: (a), (b) and (c)** represent the residue-level RMSF profiles of TAR RNA in apo vs complex state for BIV, HIV-1 and HIV-2 respectively.

**Characterizing post-Tat bound loop dynamics of HIV-2 and HIV-1 TAR by computing intra-chain RNA base-stacking probability:**

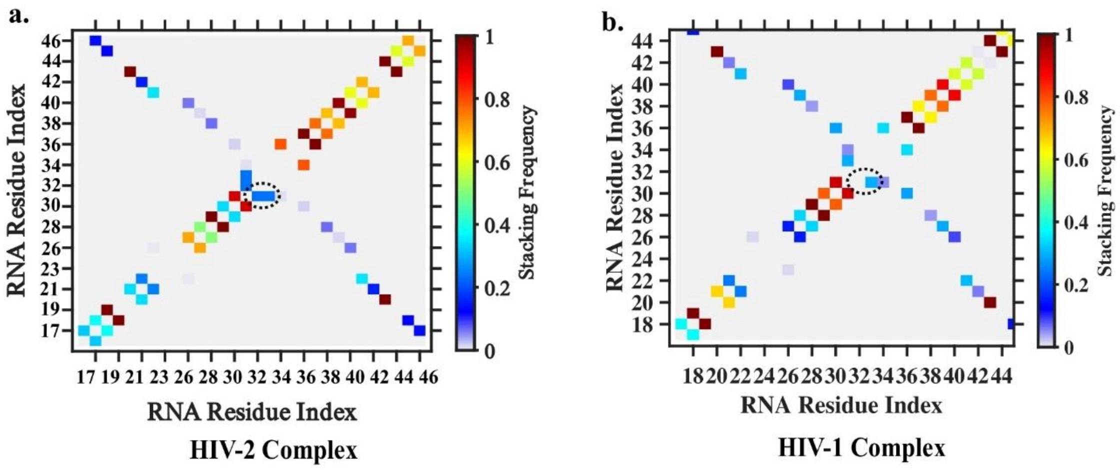


**Figure S10: Base stacking interaction comparison in complex TAR RNAs following Barnaba**(3) **definition: (a)** and **(b)** represent base stacking interaction along with the frequency of occurrence in HIV-2 and HIV-1complex TAR RNAs respectively. The existence of U31-G32 stacking interaction in loop region is circled with dotted line.

**Supporting Tables**

**Table S1: Quantification of water and ionic species around the system**

| System | No. of Water molecule | No. of Sodium ion | No. of Chloride ion |
|---|---|---|---|
| BIV TAR RNA | 56308 | 116 | 89 |
| BIV TAR-Tat Complex | 55974 | 116 | 97 |
| HIV-1 TAR RNA | 56022 | 132 | 104 |
| HIV-1 TAR-Tat Complex | 55913 | 124 | 104 |
| HIV-2 TAR RNA | 56003 | 133 | 104 |
| HIV-2 TAR-Tat Complex | 55920 | 125 | 104 |

**Table S2: Quantification of bulk water and ionic species around the system**

| System | No. of bulk Water molecule | No. of bulk Sodium ion | No. of bulk Chloride ion |
|---|---|---|---|
| BIV TAR RNA | 50306.03 | 88.35 | 83.11 |
| BIV TAR-Tat Complex | 49911.18 | 94.24 | 90 |
| HIV-1 TAR RNA | 49876.70 | 100.86 | 96.37 |
| HIV-1 TAR-Tat Complex | 49963.09 | 100.76 | 96.57 |
| HIV-2 TAR RNA | 49868.86 | 101.11 | 96.52 |
| HIV-2 TAR-Tat Complex | 49688.44 | 111.93 | 107.27 |

**Table S3: Assessment of corrected concentration of ions and preferential interaction coefficient**

| System | Ions | Rough concentration (mM) | Corrected Concentration (mM) | Preferential interaction coefficient |
|---|---|---|---|---|
| BIV TAR RNA | Na+ | 97.49 | 94.51 | 20.13 |
| | Cl- | 91.71 | 94.51 | -6.87 |

| | | | | |
|---|---|---|---|---|
| BIV TAR-Tat Complex | Na+<br>Cl- | 104.81<br>100.1 | 102.4<br>102.4 | 12.75<br>-6.25 |
| HIV-1 TAR RNA | Na+<br>Cl- | 112.25<br>107.25 | 109.69<br>109.69 | 21.29<br>-6.70 |
| HIV-1 TAR-Tat Complex | Na+<br>Cl- | 111.94<br>107.29 | 109.56<br>109.56 | 13.64<br>-6.35 |
| HIV-2 TAR RNA | Na+<br>Cl- | 112.55<br>107.44 | 109.93<br>109.93 | 22.09<br>-6.91 |
| HIV-2 TAR-Tat Complex | Na+<br>Cl- | 111.93<br>107.27 | 109.55<br>109.55 | 14.64<br>-6.36 |

**Table S4: Table for accounting total simulation in BIV case:**

| | |
|---|---|
| **Conventional MD for Apo** | Two independent 1 µs unbiased simulations = 2 µs |
| **Conventional MD for Complex** | Two independent 1 µs unbiased simulations = 2 µs |
| **SMD (COM pulling)** | After 1µs MD run equilibration: (1*50ns) = 50ns<br>After 1ns MD run equilibration: (1*50ns) = 50ns<br>After 500ns MD run equilibration: (5*40 ns) = 200ns |
| **Umbrella Windows** | 1µs MD run equilibration followed by SMD through loop: (40 windows*20ns) = 800ns<br>1ns MD run equilibration followed by SMD through loop: (30 windows*10ns) = 300ns<br>1ns MD run equilibration followed by SMD through loop: (51 windows*10ns) = 510ns |
| **Umbrella windows at high temperature (350K)** | 1ns MD run equilibration followed by SMD through loop: (30 windows*10ns) = 300ns<br>1ns MD run equilibration followed by SMD through loop: (51 windows*10ns) = 510ns |
| **Total** | **6720ns** |

**Table S5: Table for accounting total simulation in HIV-2:**

| | |
|---|---|
| **Conventional MD for Apo** | Two independent 1 µs unbiased simulations = 2 µs |
| **Conventional MD for Complex** | Two independent 1 µs unbiased simulations = 2 µs |
| **Conventional MD for Apo with $A_{35}$ positionally restrained** | Two independent 200ns long simulations = 400ns |
| **SMD (COM pulling)** | After 1µs MD run equilibration: (10*40ns) = 400ns<br>After 1ns MD run equilibration: (12*40ns) = 480ns<br>After 800ns MD run equilibration: (10*40 ns) = 400ns |
| **$A_{35}$ Position restrained directional SMD** | After 1ns MD run equilibration: 40ns |
| **RNA Position restrained SMD** | After 1ns MD run equilibration: 50ns<br>After 1µs MD run equilibration: (5*40ns) = 200ns |
| **Umbrella Windows** | 1µs MD run equilibration followed by SMD through loop: (42 windows*20ns) = 840ns<br>1µs MD run equilibration followed by SMD through terminal: (46 windows*20ns) = 920ns<br>1ns MD run equilibration (12nm box) followed by SMD through loop: (43 windows*10ns) = 430ns<br>1ns MD run equilibration followed by SMD through terminal: (42 windows*10ns) = 420ns<br>1ns MD run equilibration followed by SMD in between loop and bulge through horizontal space: (39windows*10ns) = 390ns<br>1ns MD run equilibration followed by SMD (while keeping RNA positionally restrained) in between loop and bulge through horizontal space: (49windows*10ns) = 490ns<br>800ns MD run equilibration followed by SMD through loop: (41 windows*10ns) = 410ns |
| **Umbrella windows at high temperature (350K)** | 1ns MD run equilibration followed by SMD through loop: (43 windows*10ns) = 430ns |

| | |
|---|---|
| **$A_{35}$ Position restrained umbrella windows** | 1ns MD run equilibration followed by directional SMD through loop: (43 windows*10ns) = 430ns |
| **Total** | **10730ns** |

**Table S6: Table for accounting total simulation in HIV-1:**

| | |
|---|---|
| **Conventional MD for Apo** | 1µs unbiased simulation = 1 µs |
| **Conventional MD for Complex** | 1.355µs unbiased simulation= 1.355 µs |
| **SMD (COM pulling)** | After 1µs MD run equilibration: (4*50ns) = 200ns |
| **Umbrella Windows** | 1µs MD run equilibration followed by SMD through loop: (42 windows*10ns) = 420ns<br>1µs MD run equilibration followed by SMD through loop: (41 windows*10ns) = 410ns |
| **Total** | **3385ns** |
